# vCOMBAT: a Novel Tool to Create and Visualize a COmputational Model of Bacterial Antibiotic Target-binding

**DOI:** 10.1101/2020.08.05.236711

**Authors:** Vi Ngoc-Nha Tran, Alireza Shams, Sinan Ascioglu, Antal Martinecz, Jingyi Liang, Fabrizio Clarelli, Rafal Mostowy, Ted Cohen, Pia Abel zur Wiesch

## Abstract

**Motivation:** As antibiotic resistance creates a significant global health threat, we need not only to accelerate the development of novel antibiotics but also to develop better treatment strategies using existing drugs to improve their efficacy and prevent the selection of further resistance. We require new tools to rationally design dosing regimens to from data collected in early phases of antibiotic and dosing development. Mathematical models such as mechanistic pharmacodynamic drug-target binding explain mechanistic details of how the given drug concentration affects its targeted bacteria. However, there are no available tools in the literature that allows non-quantitative scientists to develop computational models to simulate antibiotic-target binding and its effects on bacteria.

**Results:** In this work, we have devised an extension of a mechanistic binding-kinetic model to incorporate clinical drug concentration data. Based on the extended model, we develop a novel and interactive web-based tool that allows non-quantitative scientists to create and visualize their own computational models of bacterial antibiotic target-binding based on their considered drugs and bacteria. We also demonstrate how Rifampicin affects bacterial populations of Tuberculosis (TB) bacteria using our vCOMBAT tool.

**Availability:** vCOMBAT online tool is publicly available at https://combat-bacteria.org/.

## 1 Introduction

As antibiotic resistance poses a substantial worldwide health threat [1], leading academics have recently declared that we stand at the precipice of the post-antibiotic era [2]. To circumvent resistence, we need to limit inappropriate prescribing of existing drugs and also accelerate the development of novel antibiotics. Moreover, there is also a clear need to develop better treatment strategies using existing drugs to improve their efficacy and prevent the selection of further resistance.

Even though antibiotics have been used since 1944, we are not yet able to predict how antibiotic concentration affects bacteria. That leads to our inability to design rational treatment strategies using existing drugs. That is illustrated by the fact that substantial treatment improvements have been made solely based on expert opinion even after decades of clinical practice [3, 4, 5, 6, 7].

Currently, most dosing recommendations are based on the selection of the best regiments during a series of trial-and-error experiments. Many candidate drug regimens fail during this testing process, and for those candidates that do succeed, the best regimen may be missed. This costly and long trial-and-error approach may also slow down the development of new antibiotics and limits the opportunities for dosing improvement of existing drugs [8]. The design of rationale dosing of new combination regimens using multiple drugs is even more complex. The nature of the drug-drug interaction may change depending on drug concentration and therefore, antibiotic synergy and antagonism cannot usually be predicted [9]. Furthermore, differences in the pharmacokinetic and pharmacodynamic profiles of drugs used in combination regimens can promote the selection of resistance during multi-drug treatment [10, 11].

We require new tools to rationally design dosing regimens that maximize the efficacy of existing antibiotics and to shorten the development process for new antibiotics [12]. The development of models that can guide the selection of optimal dosing strategies from data collected in early phases of antibiotic development (e.g. drug-target binding and transmembrane permeability, bacteriostatic and bactericidal action of living bacteria) could accelerate the drug development process and dosing design process [13]. Computational models and tools that predict relapse from pre-clinical and early clinical data would be immensely demanded [14, 15].

Mathematical models such as mechanistic pharmacodynamic drug-target binding [16] explain mechanistic details of how the given drug concentration affects its targeted bacteria. In the mechanistic models, each living bacterium has n target molecules. The models classify living bacteria into different compartments based on the number of bound target molecules [17]. They also incorporate both bacteriostatic and bactericidal action of living bacteria into their simulations. While such models have gained traction in the last years, there are no available tools to implement those models for scientists who are not experts in mathematical modelling. Developing these computational models to simulate the mechanism of drug-target binding requires both complex modeling and programming process. For healthcare providers and scientists with a non-quantitative background, creating such mathematical models for their considered drugs and bacteria is a challenging and time-consuming task.

In this work, we have devised an extension of the mechanistic binding-kinetic model that simulates the process of bacterial antibiotic target-binding. The extended model allows the incorporation of clinical drug concentration data to the original mechanistic model [17] in order to understand the effect of drug-target binding in vivo. Based on the extended model, we have developed an interactive web-based tool, namely vCOMBAT, to allow non-quantitative scientists to create and visualize their own computational models of bacterial antibiotic target-binding. In contrast to our previously developed COMBAT modeling framework [18], this tool allows to incorporate antibiotic time-concentration profiles measured in patients. The tool can inform optimal dosing strategies based on antibiotic and bacteria data provided by the users. We also demonstrate how Rifampicin affects bacterial populations of Tuberculosis (TB) bacteria using our vCOMBAT tool.

## 2 Method and implementation

### 2.1 Mathematical models of drug binding kinetics

The web-based tool is built as an extension of the classic reaction kinetics model [17], where a bacterium has *n* target molecules binding to the antibiotic molecules. Depending on the number of bound target molecules *x* out of *n* target molecules in a bacterium, bacteria are classified into *n* + 1 compartments *B*_*x*_(where *x* is from 0 to *n*). Living bacteria also replicate and die at a rate as functions of the bound targets *x*. When a bacterium duplicates, it results in two bacteria with two times of the number of target molecules in two daughter cells. However, the number of bound target molecules *x* in the mother cell remains constant and is distributed into the two daughter cells. The distributions are calculated based on a hypergeometric distribution function.

The model is implemented as a system of ordinary differential equations as Equation 1:

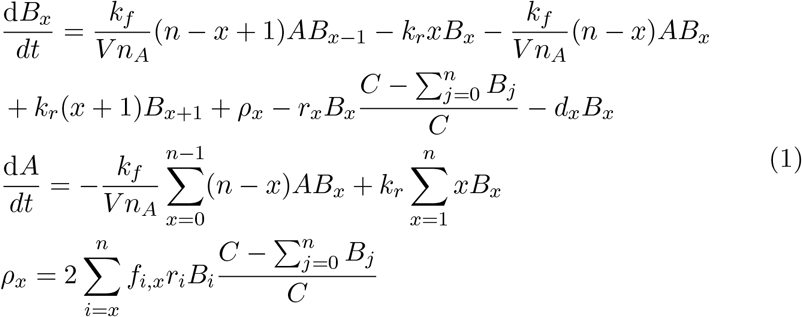

where *B*_*x*_ is the bacteria population with *x* bound targets; *n* is the number of targets per bacterium; *k*_*f*_ is the binding rate; *k*_*r*_ is the unbinding rate; *V* = *e*^*−15*^[*L/bacterialcell*] is the average intracellular volume; and *n_A_* = 6 *× e*^23^ is the Avogadro number; *C* is the carrying capacity of total bacterial population; *A* is the drug concentration; *ρ*_*x*_ is the total rate with which replication creates new bacteria with *x* bound target; *r*_*x*_ and *d*_*x*_ are the replication and death rate of bacteria with *x* bound targets, respectively; *f*_*i,x*_ is the hyper-geometric distribution function.

We develop vCOMBAT as an extension of the classic reaction kinetics model [17]. In our extension of the model, instead of calculating antibiotic concentration *A* from Equation 1, users can supply their own measured concentration data to the model. Antibiotic concentrations are measured in different time points separated by a time interval (e.g., every hour or every day). In order to incorporate the external concentration data in to the original model, the concentration values *A*(*t*) at time *t* are calculated by a linear function of the two measured concentration data points *A*(*t*_1_) and *A*(*t*_2_), where *t*_1_ *< t < t*_2_.

### 2.2 Rifampicin test case

TB is currently the bacterial infection with the highest number of infections in the world. Even though antibiotics drugs to treat TB are used for many decades, the treatment success rate is low. Understanding how anti-tuberculosis drugs affect the total bacterial population in TB patients helps to guide the design of dosing strategies. Rifampicin is one of the most effective antibiotics to treat TB due to its safety and tolerability of its high-dose treatment and its low production cost [19]. There are currently several clinical trials on assessing increasing the doses on rifampicin and, therefore, it is a huge interest to model Rifampicin actions on TB [20].

The pharmacokinetic-pharmacodynamic model is intended to capture and simulate the decrease in the number of bacteria in the cavity walls in the lungs of the tuberculosis patients in response to rifampicin exposure. Table 1 summarizes the parameter values of Rifampicin and TB bacteria used in Equations 1.

**Table 1:**
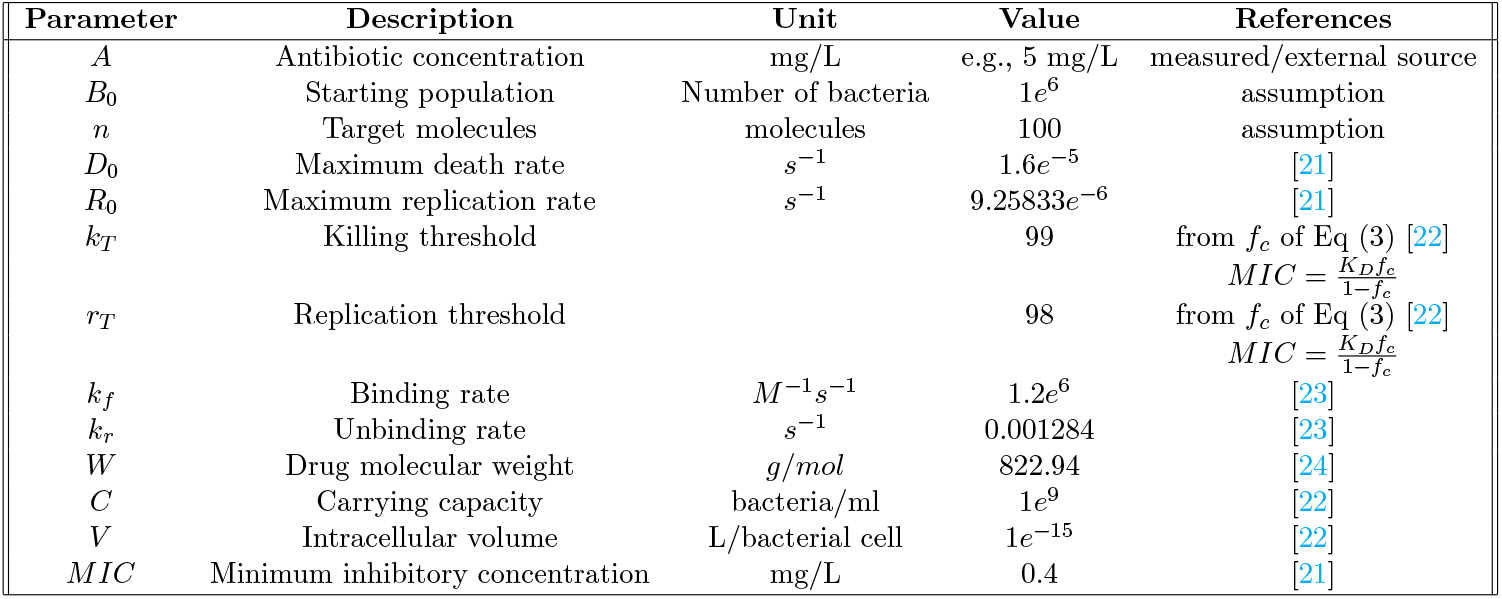
Model-parameter values when using Rifampicin in TB patients. These values are identified from the literature.

### 2.3 Model implementations

In order to make the vCOMBAT tool accessible to on-line users, it is important to have a response time (i.e., computation runtime of the model to produce results) as fast as possible. Choosing a high-performance math library for numerical computation to solve our ODE system is one of the solutions to enhance its time performance. The original model was built in R environment because R provides a vast amount of supported statistical tools and packages which makes it straightforward to program mathematical models [25]. However, R is also known for its low performance compared to other programming languages [26]. To provide high performance and short computational time, GNU Scientific Library (GSL) [27] is chosen as a numeric software package to solve our large ODE system described in Section 2.1.

We implement our original model and vCOMBAT - the extended model in C environment using GSL library and evaluate the results by comparing the simulated outputs of the original model and vCOMBAT model. Then, we conduct experiments to analyze the performance of the models.

## 3 Results

### 3.1 Model estimation of Rifampicin

In this section, we demonstrate the use of the vCOMBAT tool for antibiotic Rifampicin treatment in TB patients. We have the vCOMBAT model parameters from Table 1 and the antibiotic concentration over time from the published compartmental pharmacokinetic model from Strydom et al. [28]. The compartmental pharmacokinetic model can be used to simulate antibiotic levels in different kinds of infected tissue in the lungs of TB patients. Open cavities in the lungs are considered to be the source of the sputum which is often measured in tuberculosis clinical trials [29, 13]. Therefore, we have chosen to model the antibiotic concentrations in the tissue of cavity walls. We simplify the compartmental model to our requirement by using only absorption, plasma, and tissue compartments without using a compartment chain for the absorption as equations 2:

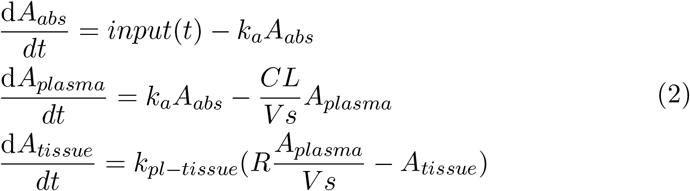

The parameters of Equation 2 are published by Strydom et al. [6], where *input*(*t*) is the input function, to be able to implement daily doses of drugs into the absorption compartment; *A*_*abs*_, *A*_*tissue*_, and *A*_*plasma*_ are antibiotic concentrations in the absorption, plasma and tissue compartments, respectively; *k*_*a*_ = 1.55[*h*^*−1*^] is the absorption rate of the drug from absorption compartment; *CL* = 5.72[*L/h*] is the clearance rate of the drug from the plasma compartment; *V s* = 52.3[*L*] is the volume of distribution in liters; *R* = 0.614 is the penetration coefficient into the tissue (cavity wall); *k*_*pl−tissue*_ = 1.98[*h*^*−1*^] rate of drug moving from plasma to tissue. For the estimates, we simulated doses of Rifampicin as 10mg/kg bodyweight (standard dose) for a 60kg person [30].

We use the antibiotic concentration over time generated from the compartmental pharmacokinetic model as concentration input for the vCOMBAT model. The simulated bacteria population over time by the vCOMBAT model is presented in Figure 1. The results show that the bacteria population reduces but then relapses approximately after day two when a patient is treated with only a single dose of Rifampicin (600 mg). The results also show that with repeated doses of Rifampicin daily (i.e., 600mg every day in 4 days), the bacteria population keeps being reduced through 4 days until 0.6% of the original population.

**Figure 1:**
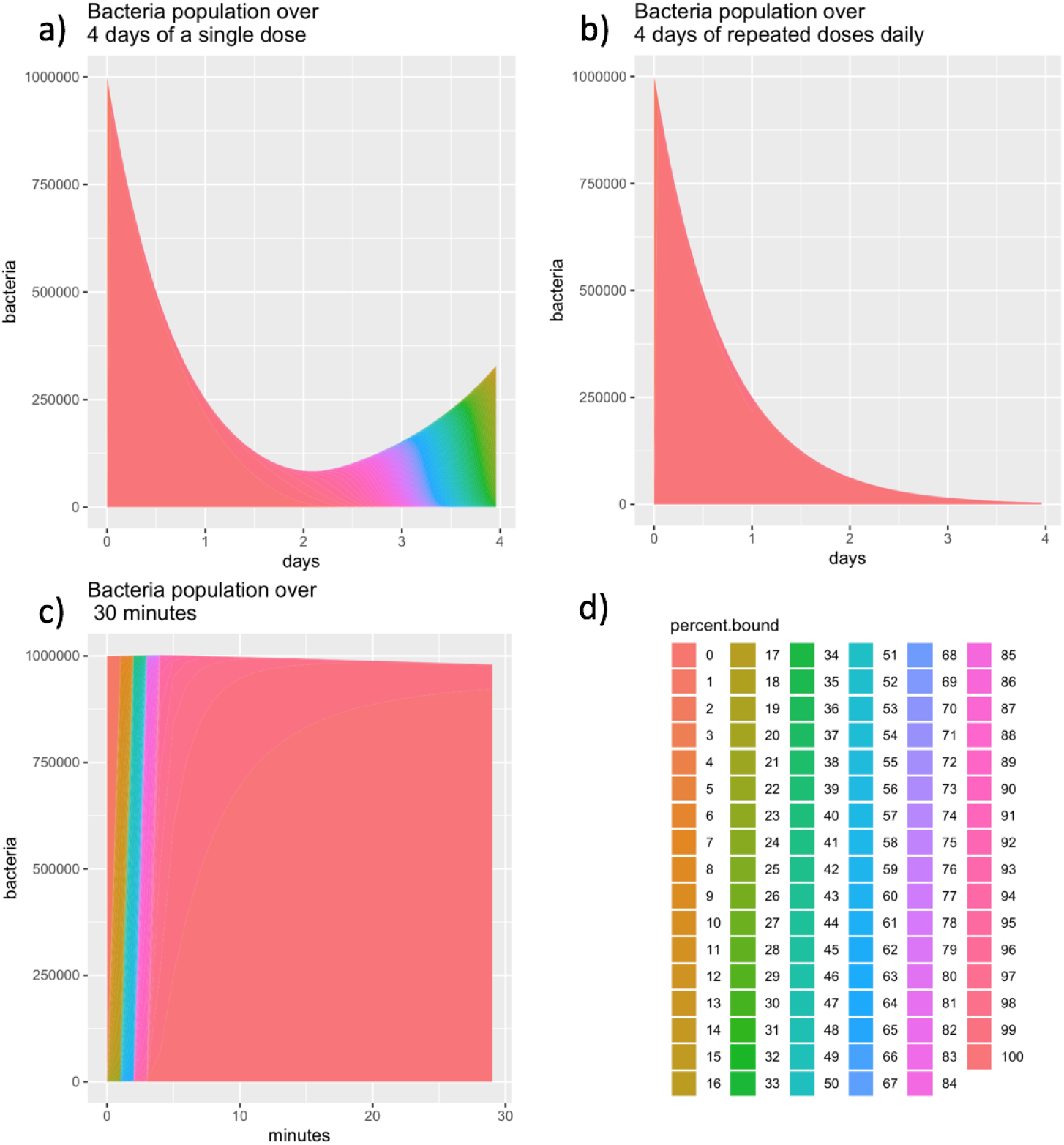
Bacteria population predicted by the extended vCOMBAT model over time. The three diagrams display the bacterial population with different simulated treatment length and dosing strategies when using Rifampicin in TB patients. The model-parameter values are taken from Table 1. The resulting graphs show that a) with a single dose of Rifampicin (600 mg), the bacteria population decreases and then regrows approximately after day two and b) with repeated doses of Rifampicin daily, the bacteria population keeps being decreased through 4 days and c) the bacteria population over the first 30 minutes of the simulated treatment for both dosing strategies in (a) and (b). The x-axis shows the simulated treatment length in hours or minutes. The y-axis shows the resulting bacteria population over the treatment time. The percent.bound legends representing the sub-populations which have from 0 to 100 percents of bound targets are depicted by different colors displayed in (d).

### 3.2 Validation with the pharmacodynamic model based on clinical data

In this section, we compare the output of our vCOMBAT model - a mechanistic pharmacodynamic model with the traditional pharmacodynamic model by Aljayyoussi et al. [31]. Mechanistic models provide a deep understanding of drug action and capture various pharmacodynamic effects [16]. Traditional models, on the other hand, are simpler but limited due to several assumptions that are likely invalid in reality. E.g., There is no cellular growth or death meaning that the total number of target molecules is constant. Traditional models are, therefore, not able to capture the pharmacodynamic effects such as post-antibiotic and inoculum effect [16].

The traditional model by Aljayyoussi et al.[31] develops the relationships of the antibiotic concentration and the net growth (elimination) rate of Mycobacterium tuberculosis bacteria exposure to Rifampicin as in Equation 3, where *A* is the antibiotic concentration in [mg/ml], *B*(*t*) represents of bacterial density over time in [*ml*^*−1*^], *r* is growth rate of bacteria in [*day*^*−1*^], *EC*_*max*_ is the maximum elimination rate in [*day^−^*^1^], and *EC*_50_ is the half-maximal effective concentration in [mg/L]. Aljayyoussi et al. [31] found the values of *EC*_*max*_ = 1.82, *EC*_50_ = 0.51, and *r* = 0.8 by fitting their clinical data into their model. With concentration *A* provided by the compartmental pharmacokinetic model [28] described in Section 3.1 and the known parameters, the bacteria population over time *B*(*t*) by the traditional model [31] is computed as Equation 3.

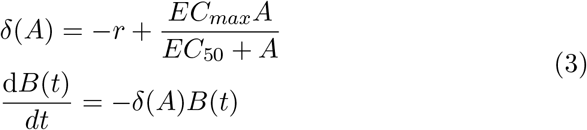

Figure 2 displays the bacteria population after treating TB patients with Rifampicin over 4 days by our mechanistic vCOMBAT model and the traditional model [31]. We notice that for a single-dose treatment (600 mg of Rifampicin) with the vCOMBAT model, the total bacteria population reduces for two days before bacteria regrow while with the traditional model, the population decreases and then increases after approximately 18 hours. This can be explained by the post-antibiotic effect [16] which the mechanistic models can capture. The post-antibiotic effect is the delay of the bacterial regrowth after bacteria are exposed to antibiotics. The bound drug-target molecules require a certain time to unbind and free the targets, as well as the drug molecules need time to leave the cell. Therefore, the vCOMBAT model in Figure 2 has the bacteria regrown later than the bacteria in the traditional model.

**Figure 2:**
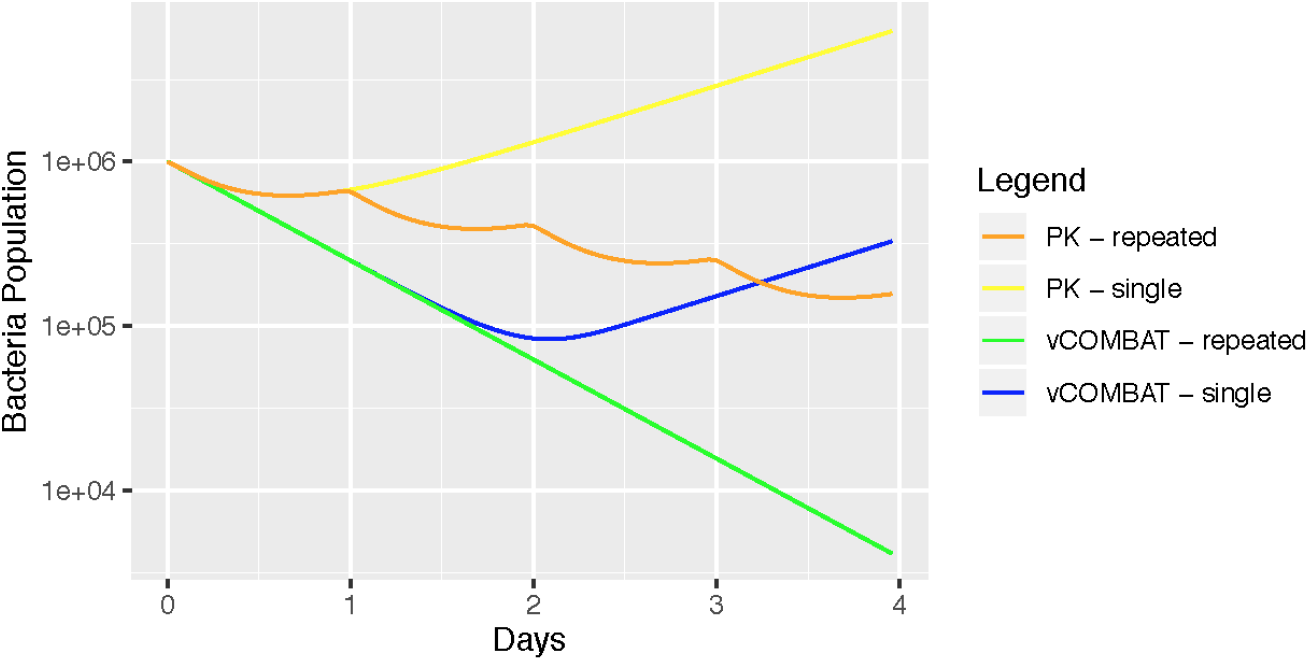
The comparison of vCOMBAT model and pharmacokinetic model regarding the bacteria population after treating TB patients with Rifampicin over 4 days. In this diagram, the x-axis shows the simulated treatment length (days). The y-axis depicts the total bacteria population throughout the treatment duration. The green and blue lines are the total bacteria population simulated by the vCOMBAT model with repeated doses and a single dose, respectively. The orange and yellow lines are the total bacteria population simulated by the pharmacokinetic model [28] with repeated doses and a single dose, respectively. The bacteria population by the vCOMBAT model has a relapse that occurred later than the population by the traditional model due to the post-antibiotic effect.

### 3.3 Model performance analysis

In this section, we analyze the performance of the original and extended vCOMBAT models. We design the three test cases for three scenarios with model parameters from Table 2. Test case 1 has no bacterial growth and death; test case 2 is a normal scenario where there is bacterial growth and death while test case 3 has a high initial antibiotic concentration which shows the effect on the bacteria subpopulations from different percentages of bound targets. To validate the results from the extended model, we compare the output of the original and the extended model for the three designed test cases. The antibiotic concentration input for the extended model is generated by the original model. In this way, we expect that the outputs of the two models are similar. Figure 3 demonstrates the effect of model parameters on the total bacteria population and bacteria population with different percentages of bound targets.

**Table 2:**
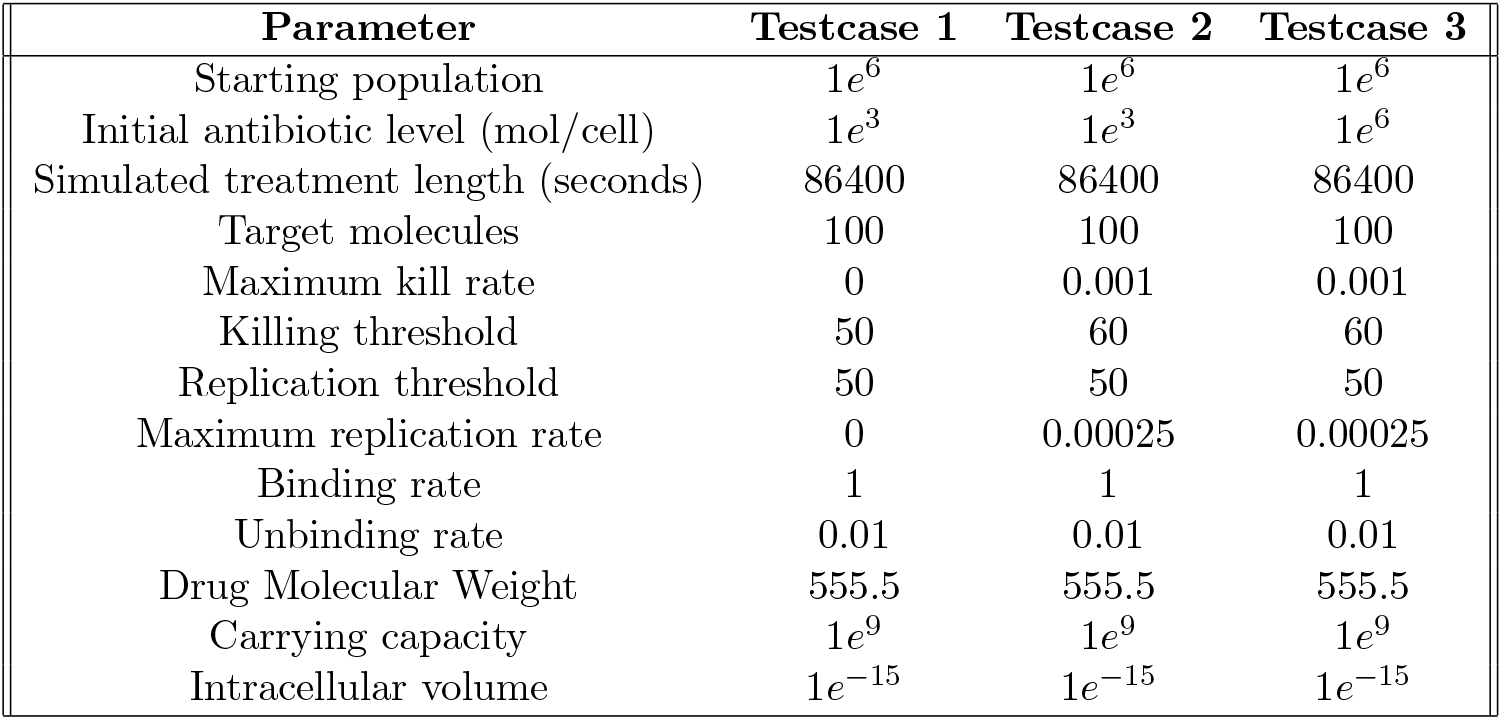
Model-parameter values of the three test cases used for model validation and performance analysis.

| Parameter | Testcase 1 | Testcase 2 | Testcase 3 |
| --- | --- | --- | --- |
| Starting population | $1e^6$ | $1e^6$ | $1e^6$ |
| Initial antibiotic level (mol/cell) | $1e^3$ | $1e^3$ | $1e^6$ |
| Simulated treatment length (seconds) | 86400 | 86400 | 86400 |
| Target molecules | 100 | 100 | 100 |
| Maximum kill rate | 0 | 0.001 | 0.001 |
| Killing threshold | 50 | 60 | 60 |
| Replication threshold | 50 | 50 | 50 |
| Maximum replication rate | 0 | 0.00025 | 0.00025 |
| Binding rate | 1 | 1 | 1 |
| Unbinding rate | 0.01 | 0.01 | 0.01 |
| Drug Molecular Weight | 555.5 | 555.5 | 555.5 |
| Carrying capacity | $1e^9$ | $1e^9$ | $1e^9$ |
| Intracellular volume | $1e^{-15}$ | $1e^{-15}$ | $1e^{-15}$ |

**Figure 3:**
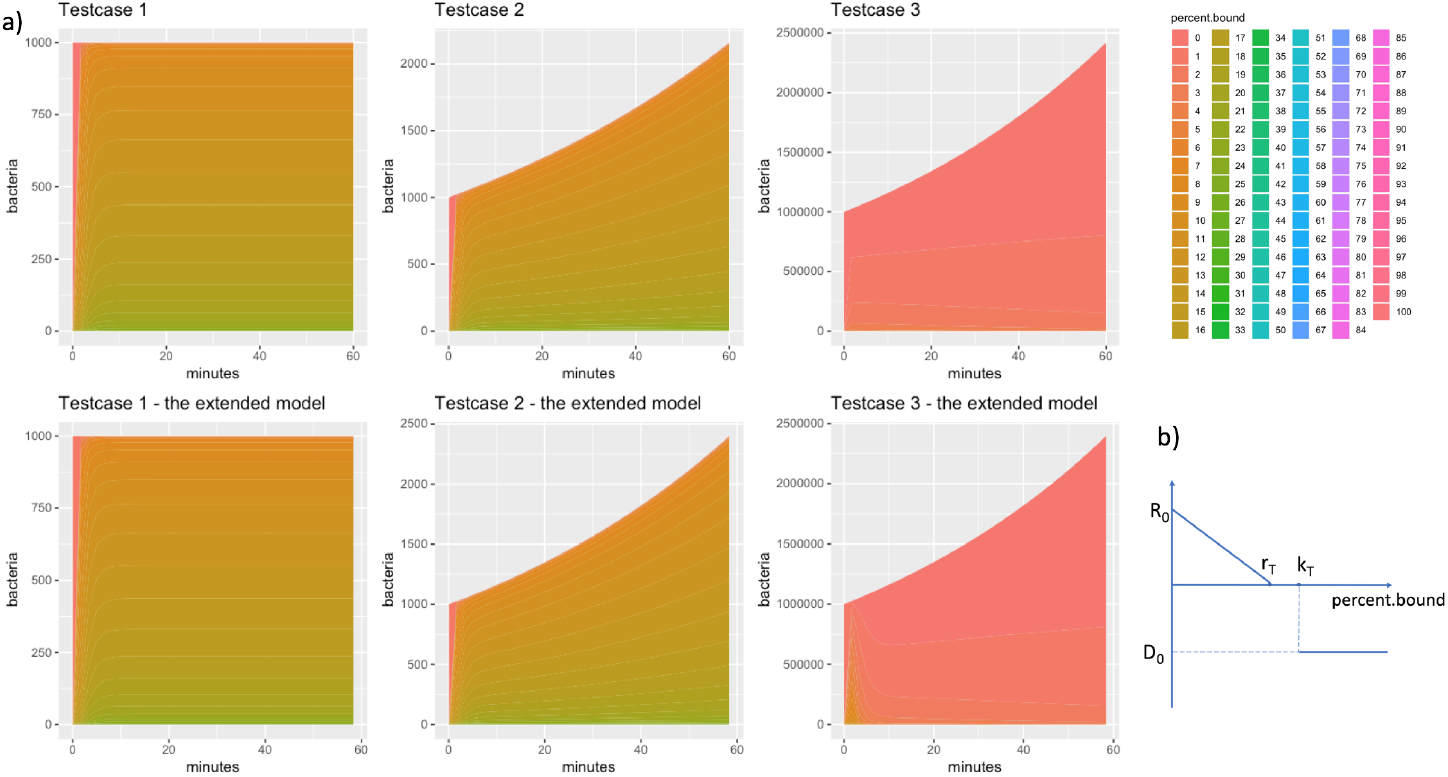
Model validation by comparing the result from the original model implemented in R and the extended model implemented in C. The three test cases were designed with different model-parameter values from Table 2 and scenarios. Test case 1 has no growth and death of bacteria. Test case 2 has the growth and death of bacteria. In test case 3, the initial dose of antibiotic is kept as the one in test case 2, but the initial number of bacteria is 1*e*^6^ instead of 1*e*^4^ as in test case 2. In (a), the x-axis shows the simulated treatment length in 60 minutes. The y-axis shows the bacteria population over the treatment length. There are 101 stacked areas representing the bacteria population which has 0 to 100 percents of bound targets. The percent.bound legends are depicted by a range of different colors. Since the external concentration input for the extended model is from the output of the original model, we expect that the two models provide similar outputs. The results show that for all three test cases, model behaviors of the original model and the extended model are similar in terms of the bacteria population and percentage bound target. In both models, the results also demonstrate the effect of model parameters such as death/growth rate, initial antibiotic level, and initial population on the final population. In test case 3, the extended model predicts an initial peak for some subpopulations due to the difference of drug-concentration profiles. I.e., the extended model is supplied with concrete values of drug concentration while the original model calculated the continuous drug concentration values at every time step. The plot (b) shows the killing curve assumed for the models, where *R*_0_, *r*_*T*_ are the maximum replication rate and replication threshold, respectively; *D*_0_, *k*_*T*_ are the death rate and killing threshold, respectively. In (b), the y-axis is the replication/death rate while the x-axis is the percentage of bound target. The more targets in the bacteria are bound, the slower rate that bacteria replicates with until replication threshold *k*_*T*_. When the percentage of bound target reaches killing threshold *k*_*T*_, the death rate becomes *D*_0_.

We analyze the performance of our extended model together with the original model in two environments: R and C. We conduct the experiments to measure computation runtime of the original model and the extended model in different environments. The original model is implemented in both R programming language and C programming language. The extended model is implemented in C environment.

#### Environmental set-up

In R environment, we used library *deSolve* to solve our ODE system and *tictoc* to measure the runtime of simulations. In C environment, we used GNU Scientific Library *GSL 2.5* to solve ODE systems. The parameters of the experiments are from Test case 2 of Table 2. Test case 2 was chosen as a typical scenario where there are growth and death of bacteria. Each experiment was run at least three times to measure the mean runtime and its variability. All experiments were conducted on an Intel platform with one Intel Core i7 processor (4 cores, 2GHz speed, and 8 GB DDR3).

#### Time performance

The performance of the original model and the extended model are illustrated in Figure 4. The experimental results show that the computational model requires significant processing time in R environment as compared to C environment. E.g., to simulate 24-hours treatment length, the computation time needs a maximum of 4252 seconds in R and a maximum 150 seconds in C. Since the computation time (runtime) is proportional to the simulated treatment length, the longer the simulated treatment length is, the longer computational time is required. The performance of the model computation is approximately 28 times faster with the conducted experiments for the extended model. By improving the performance, the model results are accessible to users in a short time. Moreover, the timely model is also beneficial in the scenario where processing algorithms require running the models with several iterations.

**Figure 4:**
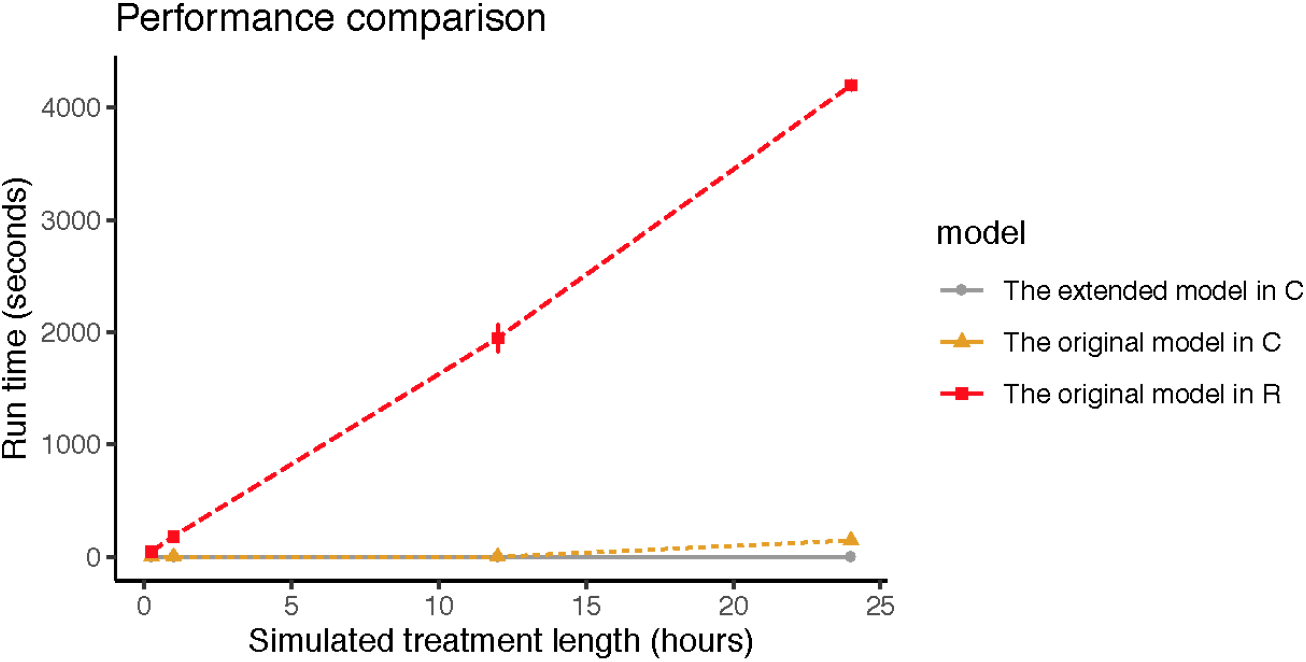
Performance comparison of the original model implemented in programming language R, the original model in C, and the extended model implemented in C. The x-axis shows the simulated treatment length in hours. The y-axis shows the runtime in seconds to complete the simulation. Each experiment (i.e., test case 2 with different simulated treatment lengths (i.e., 15 minutes, 1 hour, 12 hours, and 24 hours)) is run at least three times and their error bars represent runtime variability. The model-parameter values of experiments are from Table 2. The resulting graph shows that the computational performance of the model is improved significantly in the C environment. The extended model in C environment has the shortest computation time.

### 3.4 Implementation of vCOMBAT model into a scientific web-based tool

We develop a web-based tool to provide a user-friendly, scientific platform to create pharmacodynamic models and simulate them using our simulation software. This online tool also provides data visualization of the simulation results based on input parameter-values of the chosen drug compounds, bacteria type, and treatment length. The tool illustrates critical information such as bacteria population, drug concentration and complex bound target over treatment length to assist the design of dosing regimens. Figure 5 and Figure 6 are the sample pages visualizing the effect of a single dose of Rifampicin on a typical Tuberculosis patient over 4 days. The web tool can be freely accessed at https://combat-bacteria.org/. The tutorial providing step-by-step instructions for using the features of the interactive vCOMBAT web-based tool is in the supplementary document.The tool source code is uploaded to the git repository: https://github.com/vitrannn/vCOMBAT.

**Figure 5:**
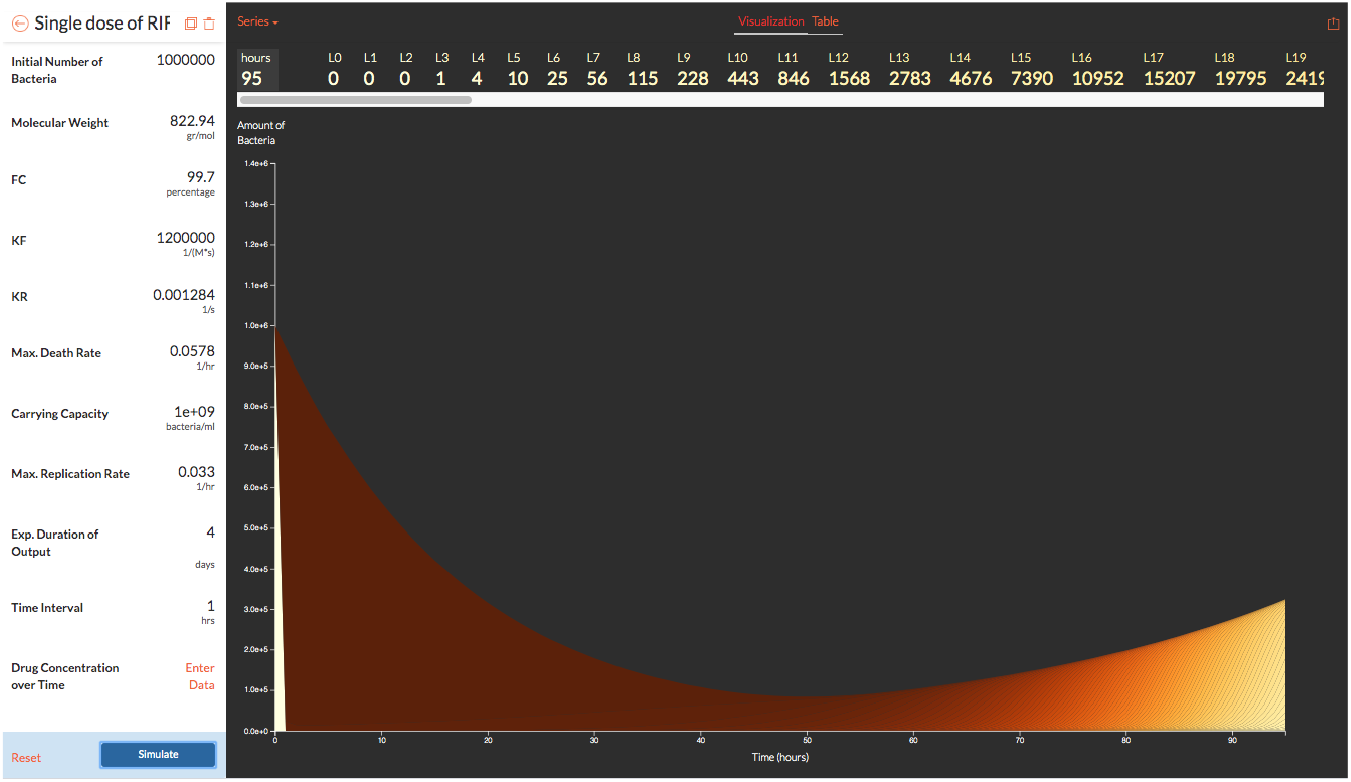
Interactive web-based tool vCOMBAT for visualizing bacteria population, antibiotic concentration, and complex bound target over simulated treatment length. This figure shows a result page of vCOMBAT displaying the bacteria population when using Rifampicin to treat TB with a single dose. The input parameters are from Table 1. The results (the graph in the right) are displayed in the logarithm scale based on model-parameter values provided by users (the panel in the left). Users can provide the desired parameter values by entering their data to the panel on the left. Users also provide measured/external antibiotic concentrations by entering data to the field “Drug Concentration over Time”. In the resulting graph, the x-axis shows the simulated treatment length in hours. The y-axis shows the resulting bacteria population in the logarithm scale over the treatment time. There are 101 stacked area in this graph representing the bacteria population *L*_*x*_ (x represents the percentage of bound targets varies from 0 to 100). The darker color depicts the higher value of x. The web-based tool also provides the output data (*L*_*x*_ values for each hour during the simulated treatment length).

**Figure 6:**
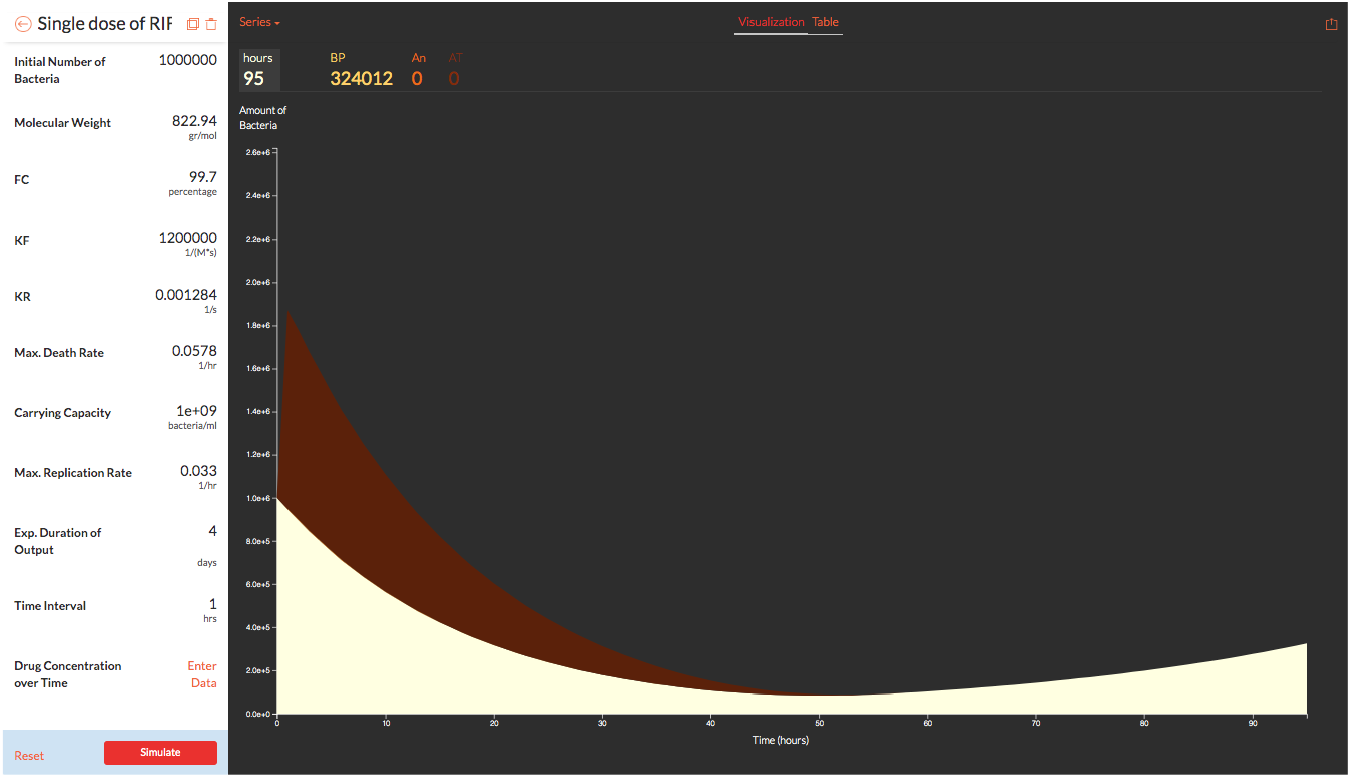
Visualizing antibiotic concentration (An) and complex bound target (AT) and the total bacteria population (BP) over simulated treatment length using the vCOMBAT web-tool. The figure shows a page of the web-based tool displaying antibiotic concentration and complex bound target when using Rifampicin to treat TB with repeated doses daily over 4 days. The results (the graph on the right) are displayed based on model-parameter values provided by users (the panel on the left). In the graph, the x-axis shows the simulated treatment length in hours. The y-axis shows the resulting complex bound target AT (the red area), the total bacteria population BP (the white area), and antibiotic concentration An (the black area, in this case, is covered by AT and BP area) over the treatment time. The user can also choose to display solely AT, BP, or An by adjusting the “Series” link in the upper-left corner of the graph.

## 4 Discussion

In the real scenario using Rifampicin to treat TB, the output of the vCOMBAT model and tool are compared with the output of the traditional pharmacodynamic model. Both models can predict the relapse from a single-dose Rifampicin at different time points. The difference is accounted for by the post-antibiotic effect. The vCOMBAT model is a mechanistic pharmacodynamic model that can capture the post-antibiotic effect where bacterial regrowth is delayed. The result from the vCOMBAT tool can aid the selection of an optimal drug dosing by informing which dosing regimens can terminate the bacterial population and clear an infection. It can also predict relapse from pre-clinical and early clinical data and therefore, shorten the development process for new antibiotics.

The vCOMBAT tool run time varies by the values of the model parameters. For the model of using Rifampicin to treat TB patients, the runtime is longer (e.g., 15 minutes of simulation for 4 days of simulated treatment length) than the sample test cases due to the values of killing and replication threshold. The combination of Rifampicin drug and TB has extreme values of killing and replication threshold (i.e., TB bacterium are killed when 99 of its 100 free target molecules are bound). That means at every time step, the ODE solver has to compute 99 sub-populations of compartments *B*_*x*_ and assure their precision at the same time. However, the tool runtime (i.e., 15 minutes) is still considerably quick given the long simulated treatment length (i.e., 4 days).

The vCOMBAT tool provides a user-friendly and scientific platform for non-quantitative scientists and healthcare providers to create and visualize their own binding kinetic models for their considered drugs and bacteria. Moreover, with a timely and interactive tool, it also opens a wide range of opportunities to further use the vCOMBAT model in practices. The model can predict the drug efficacy for a large selection of dosing regimens and guide the choice of optimal doses. It can also be integrated with machine learning techniques to automatize the process of selecting optimal dosing.

## 5 Conclusions

This work developed an extension of the mechanistic binding-kinetic model that simulates the process of bacterial antibiotic target-binding and presents the effect of drug actions on bacterial population over time. Based on the vCOMBAT model, we developed an interactive online tool that allows scientists and healthcare providers to create and visualize their own binding-kinetic models in a quick response time. We also demonstrated how the vCOMBAT tool simulates and visualizes the effect by different dosing strategies of Rifampicin on TB bacterial populations.

In the future, this work will be developed further to devise a framework to assist the process of chosing the optimal dosings. In the case where there is a wide range of possible dosings to be considered, modeling and selecting the optimal dosing from all the dosing possibilities are significantly more complex. Our ultimate aim is to make the process of selecting optimal dosing less time-consuming which is critical in improving patient well-being.

## Supporting information

vCOMBAT Tutorial

## References

[1] United Nations. http://www.un.org/pga/71/event-latest/high-level-meeting-on-antimicrobial-resistance/(2020-04-02), 2016.

[2] Mark Woolhouse and Jeremy Farrar. Policy: An intergovernmental panel on antimicrobial resistance. Nature, 509(7502):555–557, 2014.

[3] Martin J. Boeree, Andreas H. Diacon, Rodney Dawson, Kim Narunsky, Jeannine Du Bois, Amour Venter, Patrick P.J. Phillips, Stephen H. Gillespie, Timothy D. McHugh, Michael Hoelscher, Norbert Heinrich, Sunita Rehal, Dick Van Soolingen, Jakko Van Ingen, Cecile Magis-Escurra, David Burger, Georgette Plemper Van Balen, and Rob E. Aarnoutse. A dose-ranging trial to optimize the dose of rifampin in the treatment of tuberculosis. American Journal of Respiratory and Critical Care Medicine, 191(9):1058–1065, 2015.

[4] A. J. Lan, J. M. Colford, and J. M. Colford. The impact of dosing frequency on the efficacy of 10-day penicillin or amoxicillin therapy for streptococcal tonsillopharyngitis: A meta-analysis. Pediatrics, 105(2), 2000.

[5] J J Roord, B H Wolf, M M Gossens, and J L Kimpen. Prospective open randomized study comparing efficacies and safeties of a 3-day course of azithromycin and a 10-day course of erythromycin in children with community-acquired acute lower respiratory tract infections. Antimicrobial Agents and Chemotherapy, 40(12):2765–2768, 1996.

[6] A Van Deun, M A Hamid Salim, A P Kumar Das, I Bastian, and F Portaels. Results of a standardised regimen for multidrug-resistant tuberculosis in Bangladesh A standardised approach may provide a reasonable alternative to individualised treatment of MDR-TB in resource-poor settings. However, DOTS-plus programmes in resource-p. Int J Tuberc Lung Dis, 8(5):560–567, 2004.

[7] WHO. The shorter mdr-tb regimen., 2016.

[8] C. Robert Horsburgh, Clifton E. Barry, and Christoph Lange. Treatment of tuberculosis. New England Journal of Medicine, 373(22):2149–2160, 2015. PMID: 26605929.

[9] Peter Ankomah and Bruce R. Levin. Two-drug antimicrobial chemotherapy: A mathematical model and experiments with Mycobacterium marinum. PLoS Pathogens, 8(1), 2012.

[10] Qiucen Zhang, Guillaume Lambert, David Liao, Hyunsung Kim, Kristelle Robin, Chih-kuan Tung, Nader Pourmand, and Robert H. Austin. Acceleration of emergence of bacterial antibiotic resistance in connected microenvironments. Science, 333(6050):1764–1767, 2011.

[11] Christopher Dye, Brian G. Williams, Marcos A. Espinal, and Mario C. Raviglione. Erasing the world’s slow stain: Strategies to beat multidrug-resistant tuberculosis. Science, 295(5562):2042–2046, 2002.

[12] S Ragnar Norrby, Carl Erik Nord, and Roger Finch. Lack of development of new antimicrobial drugs: a potential serious threat to public health. European Society of Clinical Microbiology and Infectious Diseases (ESCMID), 2005.

[13] Geraint Davies, Martin Boeree, Dave Hermann, and Michael Hoelscher. Accelerating the transition of new tuberculosis drug combinations from phase ii to phase iii trials: New technologies and innovative designs. PLOS Medicine, 16(7):1–10, 07 2019.

[14] Rumin Zhang. Pharmacodynamics which trails are your drugs taking? Nature Chemical Biology, 11:382–383, 2015.

[15] Christian Lienhardt and Payam Nahid. Advances in clinical trial design for development of new tb treatments: A call for innovation. PLOS Medicine, 16(3):1–5, 03 2019.

[16] Fabrizio Clarelli, Jingyi Liang, Antal Martinecz, Ines Heiland, and Pia Abel zur Wiesch. Multi-scale modeling of drug binding kinetics to predict drug efficacy. Cellular and Molecular Life Sciences, 77(3):381–394, 2020.

[17] Pia Abel Zur Wiesch, Sören Abel, Spyridon Gkotzis, Paolo Ocampo, Jan Engelstädter, Trevor Hinkley, Carsten Magnus, Matthew K. Waldor, Klas Udekwu, and Ted Cohen. Classic reaction kinetics can explain complex patterns of antibiotic action. Science Translational Medicine, 7(287):1–12, 2015.

[18] F. Clarelli, A. Palmer, B. Singh, M. Storflor, S. Lauksund, T. Cohen, and P. Abel zur Wiesch S. Abel. Drug-target binding quantitatively predicts optimal antibiotic dose levels in quinolones. Plos Comp. Biol., 2020.

[19] Melanie Grobbelaar, Gail E. Louw, Samantha L. Sampson, Paul D. Helden, Peter R. Donald, and Robin M. Warren. Evolution of rifampicin treatment for tuberculosis. Infection, Genetics and Evolution, 74:103937, 2019.

[20] Martin J. Boeree, Norbert Heinrich, Rob Aarnoutse, Andreas H. Diacon, Rodney Dawson, Sunita Rehal, Gibson S. Kibiki, Gavin Churchyard, Ian Sanne, Nyanda E. Ntinginya, Lilian T. Minja, Robert D. Hunt, Salome Charalambous, Madeleine Hanekom, Hadija H. Semvua, Stellah G. Mpagama, Christina Manyama, Bariki Mtafya, Klaus Reither, Robert S. Wallis, Amour Venter, Kim Narunsky, Anka Mekota, Sonja Henne, Angela Colbers, Georgette Plemper van Balen, Stephen H. Gillespie, Patrick P.J. Phillips, and Michael Hoelscher. High-dose rifampicin, moxifloxacin, and SQ109 for treating tuberculosis: a multi-arm, multi-stage randomised controlled trial. The Lancet Infectious Diseases, 17(1):39–49, 2017.

[21] Dominique Cadosch, Pia Abel zur Wiesch, Roger Kouyos, and Sebastian Bonhoeffer. The Role of Adherence and Retreatment in De Novo Emergence of MDR-TB. PLoS Computational Biology, 12(3):1–19, 2016.

[22] Pia Abel zur Wiesch, Fabrizio Clarelli, and Ted Cohen. Using chemical reaction kinetics to predict optimal antibiotic treatment strategies. PLOS Computational Biology, 13(1):1–28, 01 2017.

[23] Walter Wehrli. Kinetic Studies of the Interaction between Rifampicin and DNA-Dependent RNA Polymerase of Escherichia coli. Biological Research, 330:325–330, 1977.

[24] Compound summary of rifampicin. https://pubchem.ncbi.nlm.nih.gov/compound/rifampicin (2019-01-15), 2019.

[25] R Core Team. R: A Language and Environment for Statistical Computing. R Foundation for Statistical Computing, Vienna, Austria, 2014.

[26] S. Borağan Aruoba and Jesús Fernández-Villaverde. A comparison of programming languages in economics. Journal of Economic Dynamics and Control, 58:265–273, June 2015.

[27] m. et al Galassi. Gnu scientific library reference manual, 2018.

[28] Natasha Strydom, Sneha V. Gupta, William S. Fox, Laura E. Via, Hyeeun Bang, Myungsun Lee, Seokyong Eum, Tae Sun Shim, Clifton E. Barry, Matthew Zimmerman, Véronique Dartois, and Radojka M. Savic. Tuberculosis drugs’ distribution and emergence of resistance in patient’s lung lesions: A mechanistic model and tool for regimen and dose optimization. PLoS Medicine, 16(4):1–26, 2019.

[29] Kelly E. Dooley, Debra Hanna, Vidya Mave, Kathleen Eisenach, and Radojka M. Savic. Advancing the development of new tuberculosis treatment regimens: The essential role of translational and clinical pharmacology and microbiology. PLOS Medicine, 16(7):1–14, 07 2019.

[30] Jakko Van Ingen, Rob E. Aarnoutse, Peter R. Donald, Andreas H. Diacon, Rodney Dawson, Georgette Plemper van Balen, Stephen H. Gillespie, and Martin J. Boeree. Why Do We Use 600 mg of Rifampicin in Tuberculosis Treatment? Clinical Infectious Diseases, 52(9):e194–e199, 05 2011.

[31] Ghaith Aljayyoussi, Victoria A. Jenkins, Raman Sharma, Alison Ardrey, Samantha Donnellan, Stephen A. Ward, and Giancarlo A. Biagini. Pharmacokinetic-Pharmacodynamic modelling of intracellular Mycobacterium tuberculosis growth and kill rates is predictive of clinical treatment duration. Nature Scientific Reports, 7(1):1–11, 2017.

