## Supplementary material for "vCOMBAT: a Novel Tool to Create and Visualize a COmputational Model of Bacterial Antibiotic Target-binding": vCOMBAT Tutorial

August 3, 2020

### 1 Introduction

The vCOMBAT online tool is hosted at https://combat-bacteria.org/. The tool can be accessed by any web browsers. Only web browser is required.

The features of vCOMBAT tool include the follows: create a drug-target binding model based on user chosen drug and bacteria in a user-friendly interface via entering model parameters, provide data visualization of simulation results, store and share models.

In this tutorial, the instructions are provided for the following tasks: create a user account, create and simulate a model by providing model parameters, visualize and interpret your output data, store and share your models.

### 2 Tutorial Tasks

#### 2.1 Homepage and description

From homepage, users can see a sample model as Figure 1 or description as Figure 2.

#### 2.2 Create a new account and sign in

**Step 1: Go to log in/sign-up page** Users click on "Join Now to Create Model" to go to sign-up page or click on "My Models" to go to log-in page as Figure 3

**Step 2: Filling the desired parameter for your model** At sign-up page, users fill in the desired username, email address and password to create an account. Users need an account in order to create their own

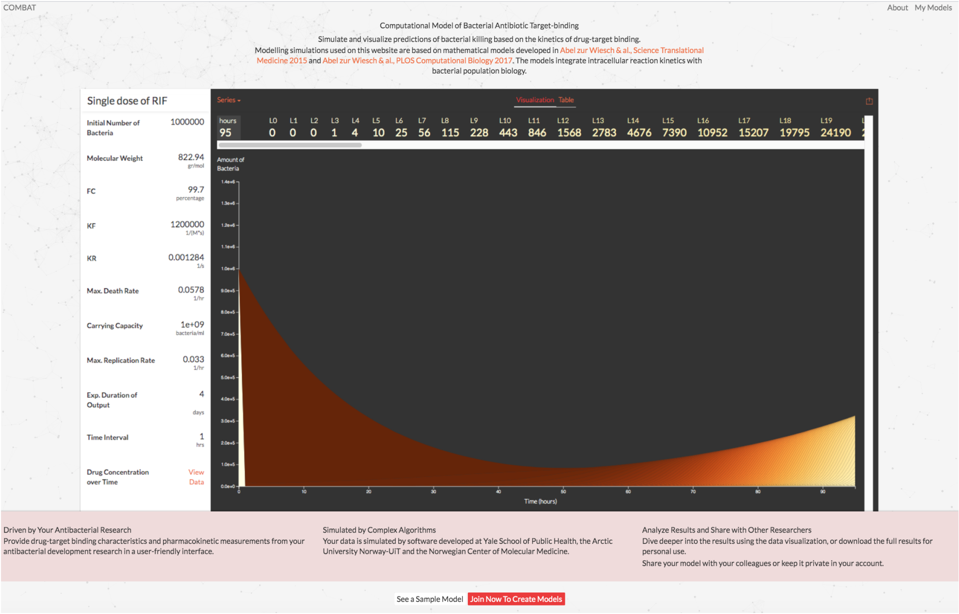


Figure 1: Homepage

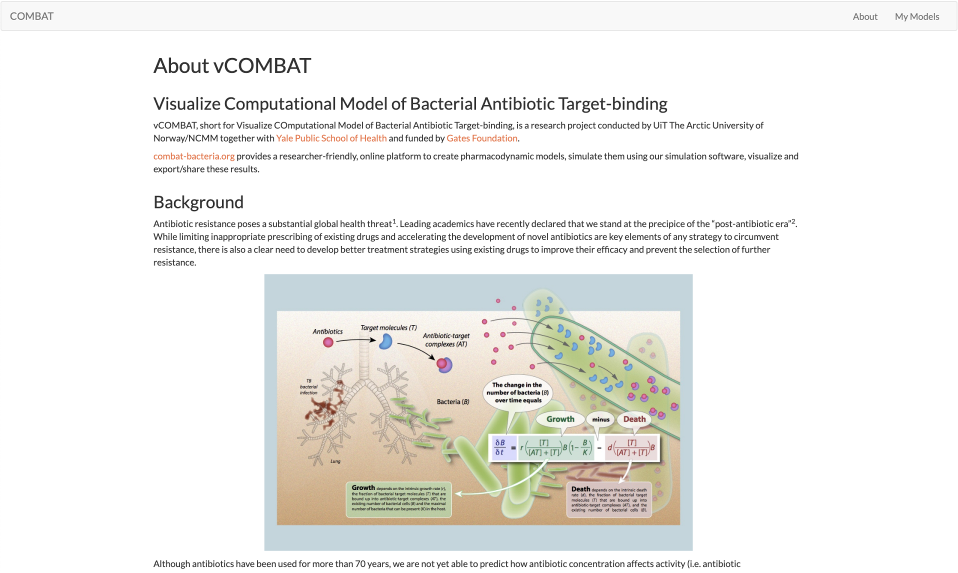


Figure 2: Description page

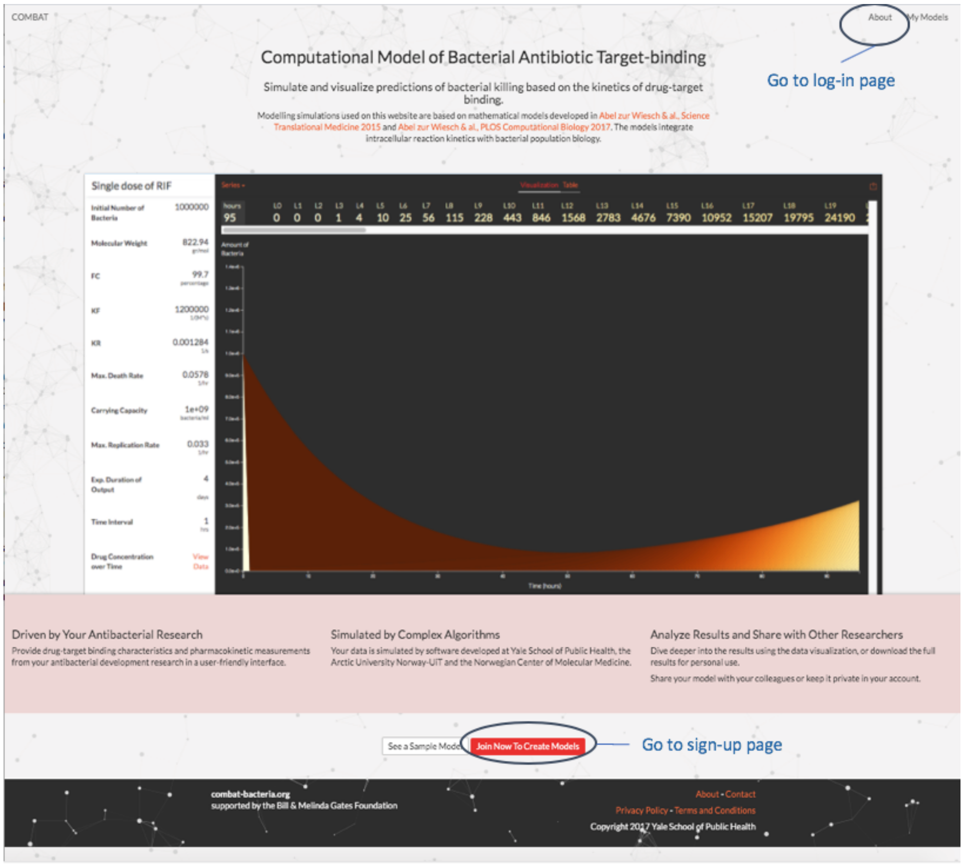


Figure 3: Go to log-in/sign-up page

models. After filling the mentioned information, users click on the checkbox "I'm not a robot" and then click "Join" as Figure 4.

**Step 3: Sign in with the created account (email and password)** Go to the log-in page and enter user email and password from the created account. Then, click "Sign in" as Figure 5.

#### 2.3 Fill in model parameters to create and simulate a model

**Step 0 After log-in to your account, click on "Create Model"** After creating a model, a model with default parameters will be assigned. The snapshot in Figure 6 shows how the process of Step 1 and 2.

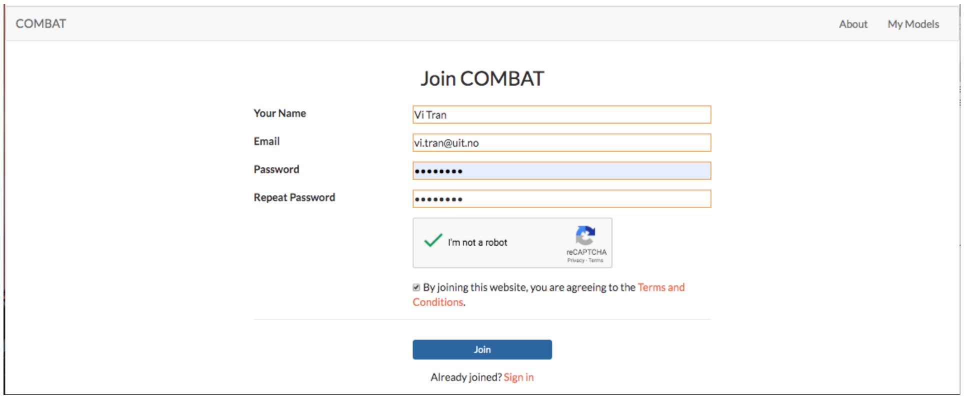


Figure 4: Sign-up window

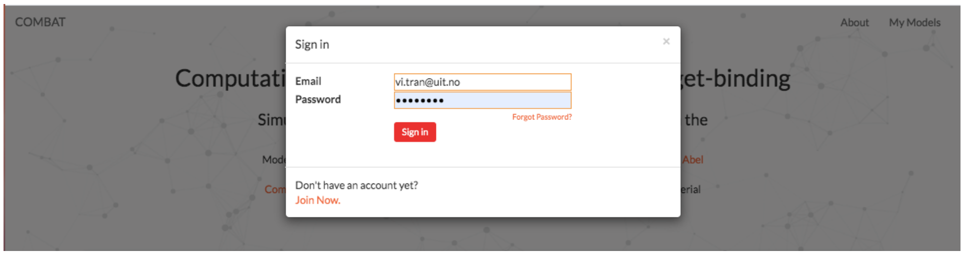


Figure 5: Log-in window

**Step 1 Fill in the desired parameters regarding treatment length, drug and bacteria** FC is computed based on MIC and KD (kf, kr) following Eq 3 $MIC = \frac{K_D f_c}{1-f_c}$ [?]. The model parameter values we used in this tutorial are as follows.

- Initial number of bacteria = 1000000
- Molecular weight = 822.94 gr/mol
- Fc = 99.7
- Kf = 1200000 $M^{(} - 1)s^{(} - 1)$
- Kr = 0.001284 $s^{(} - 1)$
- Max. death rate = 0.0578 hr-1
- Carrying capacity = 1000000000

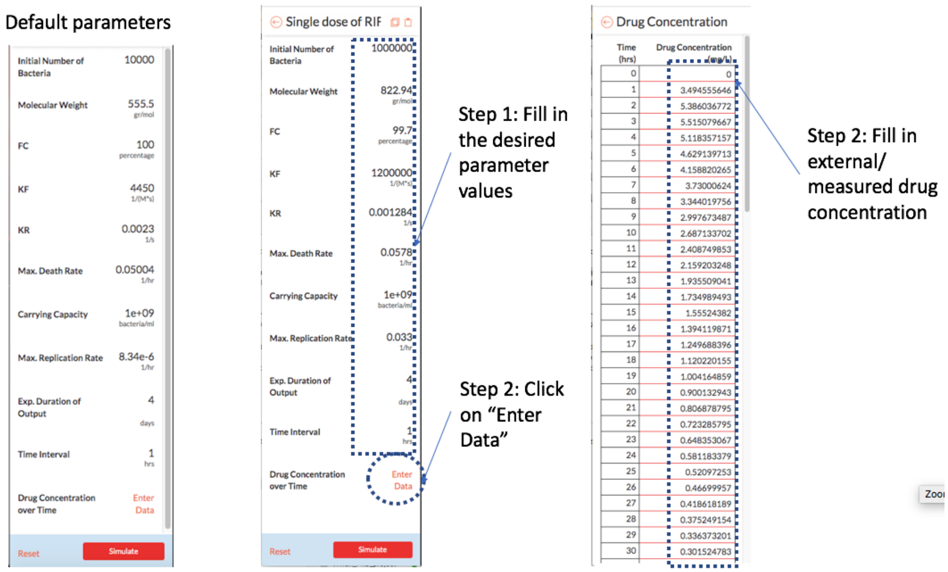


Figure 6: Filling in model parameters

- Max. replication rate = 0.0333 $h^(-1)$
- Exp. duration of output = 4 day (96 hours)
- Time interval = 1 hour

**Step 2 Fill in external/measured drug concentration over time (in mg/L) by clicking on "Enter Data"** The time interval for drug concentration is set via "time interval" textbox. The interface for entering external concentration data looks like below. The drug concentration values can be pasted directly from a column of data in an external data or excel file as Figure Figure 6 .

**Step 3 Start simulation** After entering the external drug concentrations, you can go back to the model page via the arrow near "Drug Concentration" and click on 'Simulate' button to start model simulation. It is noted that the computation time of the simulation depends not only on the treatment length but also on the values of model parameters. With the parameter set from Table 2 and a single dose of Rifampicin, the expected computation time to simulate the model for one day treatment length is approximately 6 minutes and four days is approximately 15 minutes. User can choose to have a shorter treatment length by filling ”Exp. Duration of Output” to have the simulated output in a shorter time. Figure 7 is the sample snapshot when the model is simulating.

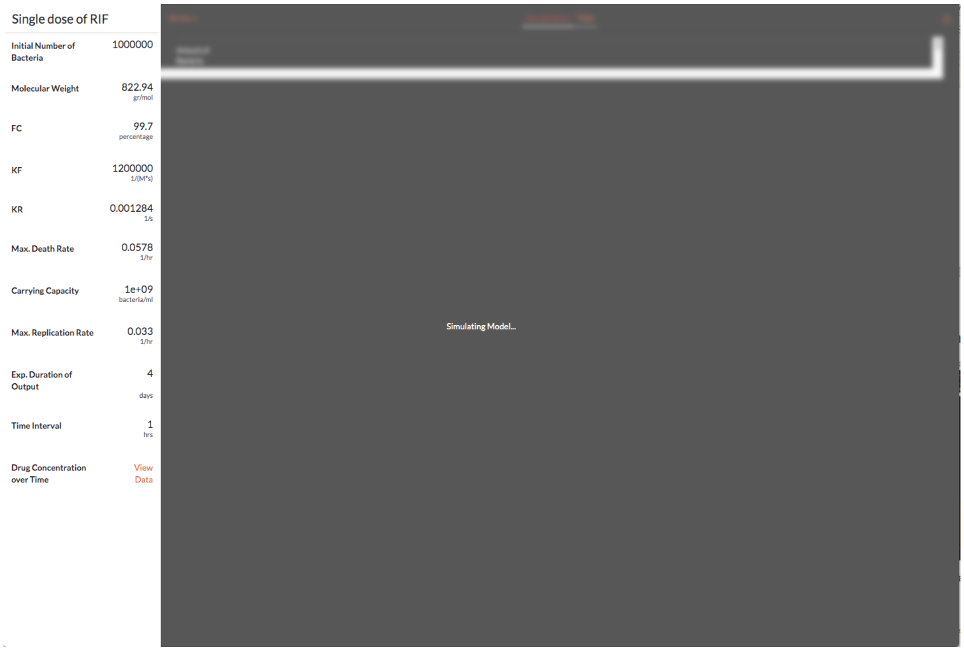


Figure 7: Simulating the model

After simulating the model from the previous task, total bacteria population and sub-population of different percentages of bound targets (L0 to L100) are displayed as Figure 8.

### 2.4 Visualize and interpret model results with graphs and data tables

Users can choose to display one of the two data series in graphs. Data series one includes BP (bacteria population), An (Antibiotic) and AT (Antibiotic-target complex). Data series two includes different bacteria populations with x percentages of bound targets Lx (where x ranges from 0 to 100).

**Step 1: Choose series one as the below figure** The sample snapshots of Step 1 and 2 are as Figure 9 and 10, respectively.

**Step 2: Choose series two as the below figure**

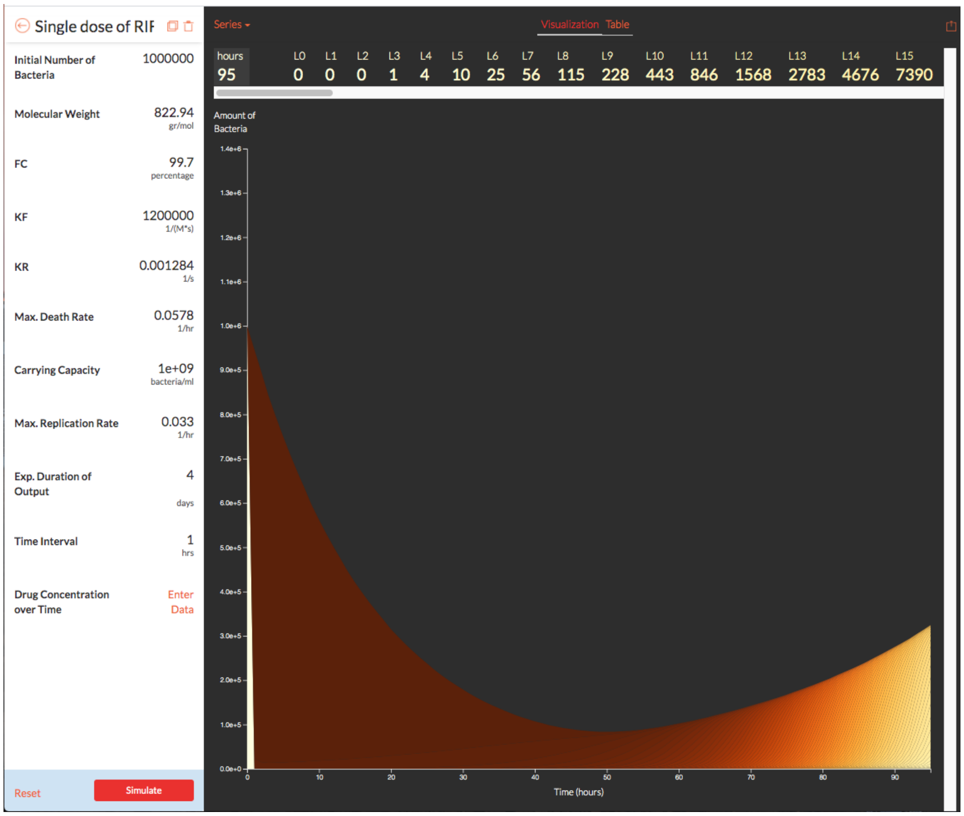


Figure 8: After simulation

**Step 3: See the series 1' data** Users can also choose to display the data values instead of graphs by using the tab "Table". Then choose the series. When choosing series 1, there are 4 columns of data in the below figure representing time interval index, Bacteria Population (BP), Antibiotic (An in mol/cell) and Complex Bound Target AT as Figure 11.

**Step 4: See the series 2' data** When choosing series 2, there are 102 columns of data in the below figure representing time interval index, L0 to L100 as Figure 12.

**Step 5: Choose specific data among the series** Users can also choose to display specific data values on each series. For example, only displaying L0 and L100 as the below diagram as Figure 13.

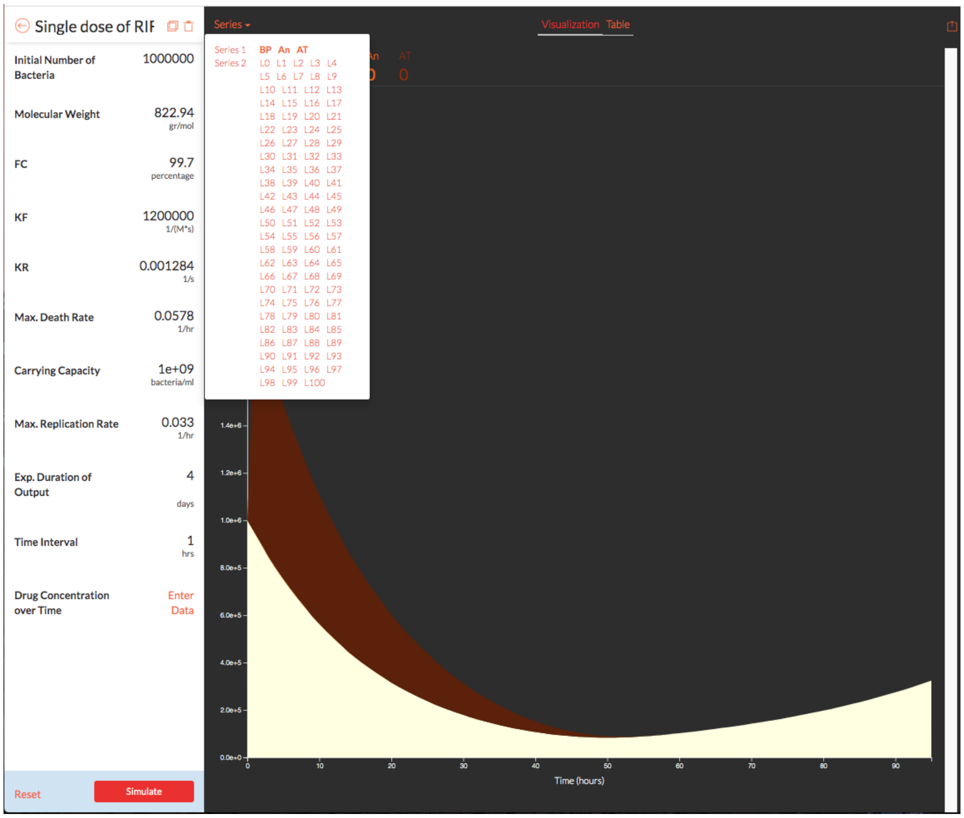


Figure 9: Choose series 1 diagram

### 2.5 Download the output data and share the model by sending link or email

**Step 1: Drop-down list** Choose the desired option from the drop-down list: Download Table, Share Link or Email Model from the dropdown list as Figure 14.

**Step 2: Share link** After choosing "Share Link", another browser tab is generated with the url link as Figure 15.

**Step 3: Email Model** After choosing "Email Model", an email is generated and the model url is included in the email text. By this point, you have gone through the features of our webtool, including create a new account, create a model and fill in desired values for model parameters, simulate a

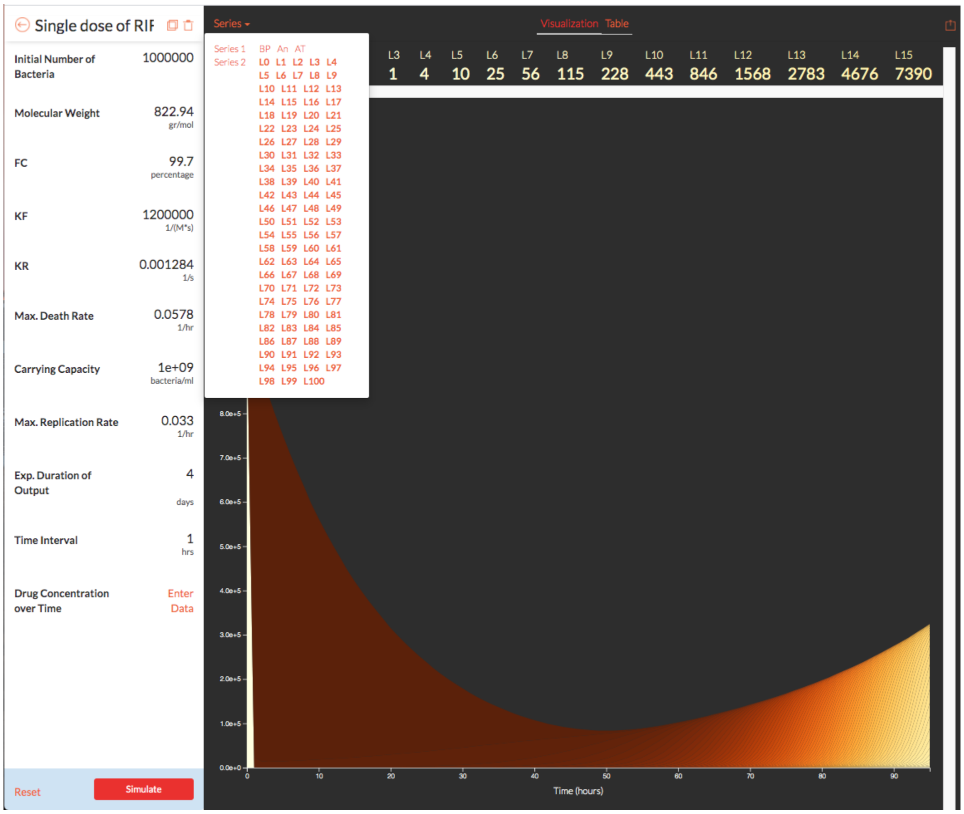


Figure 10: Choose series 2 diagram

model and visualize the output data to see how antibiotics affect the bacteria population, save and share the results and the model.

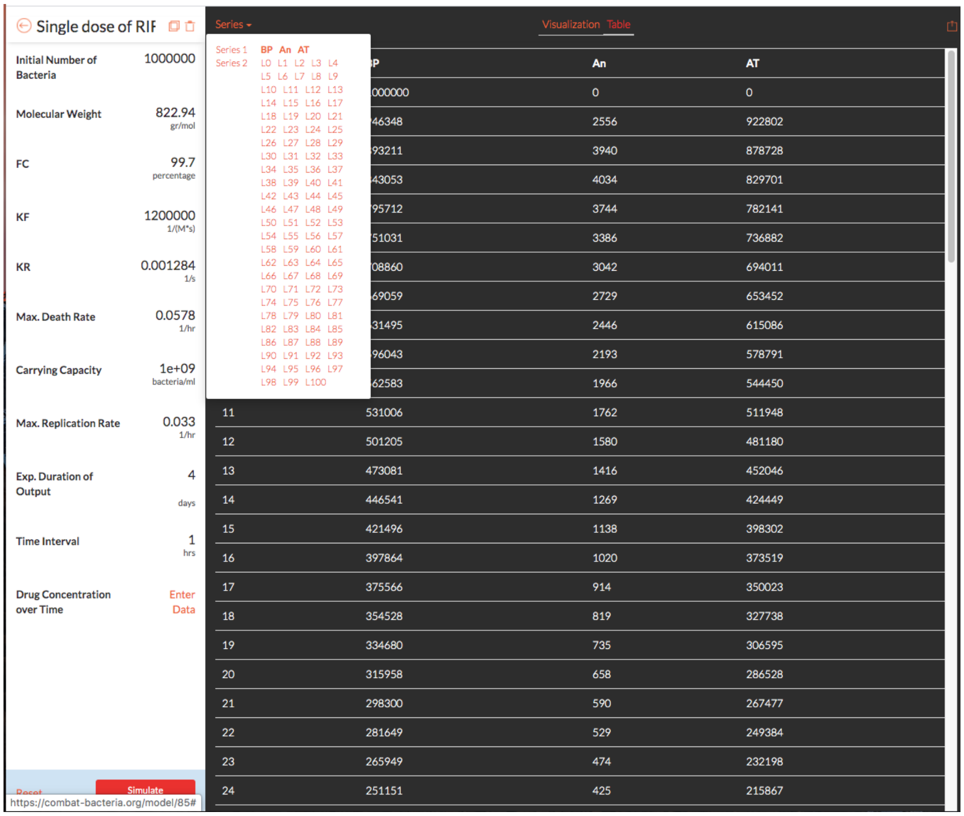

Figure 11: Choose series 1 data

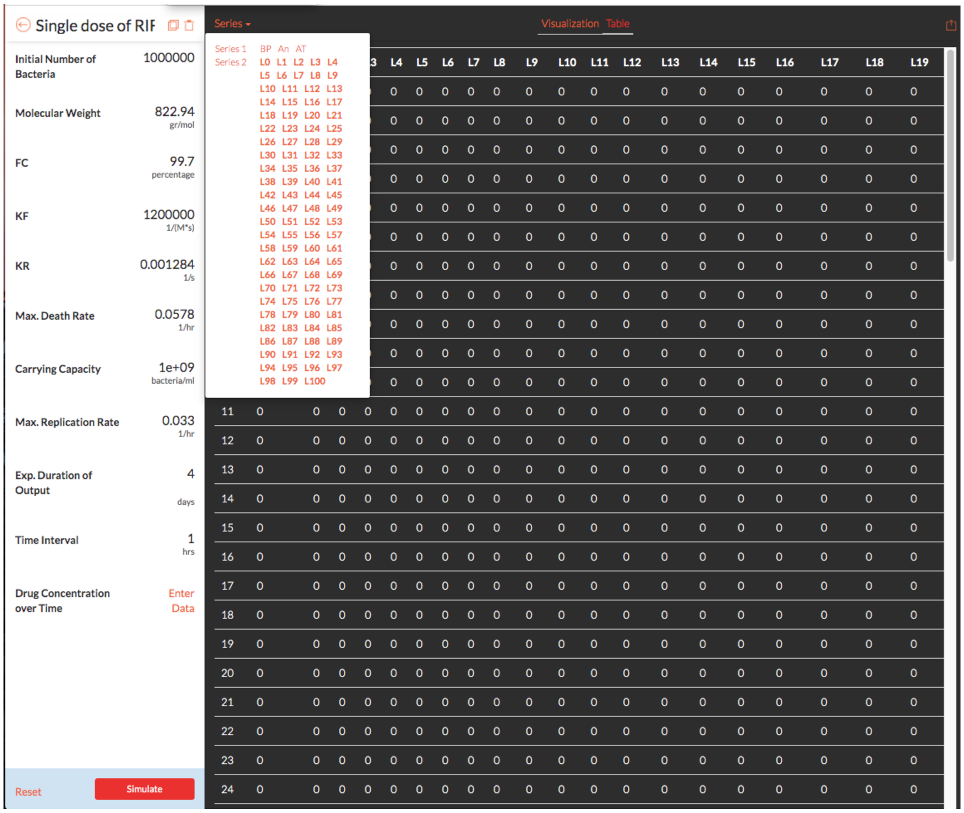

Figure 12: Choose series 2 data

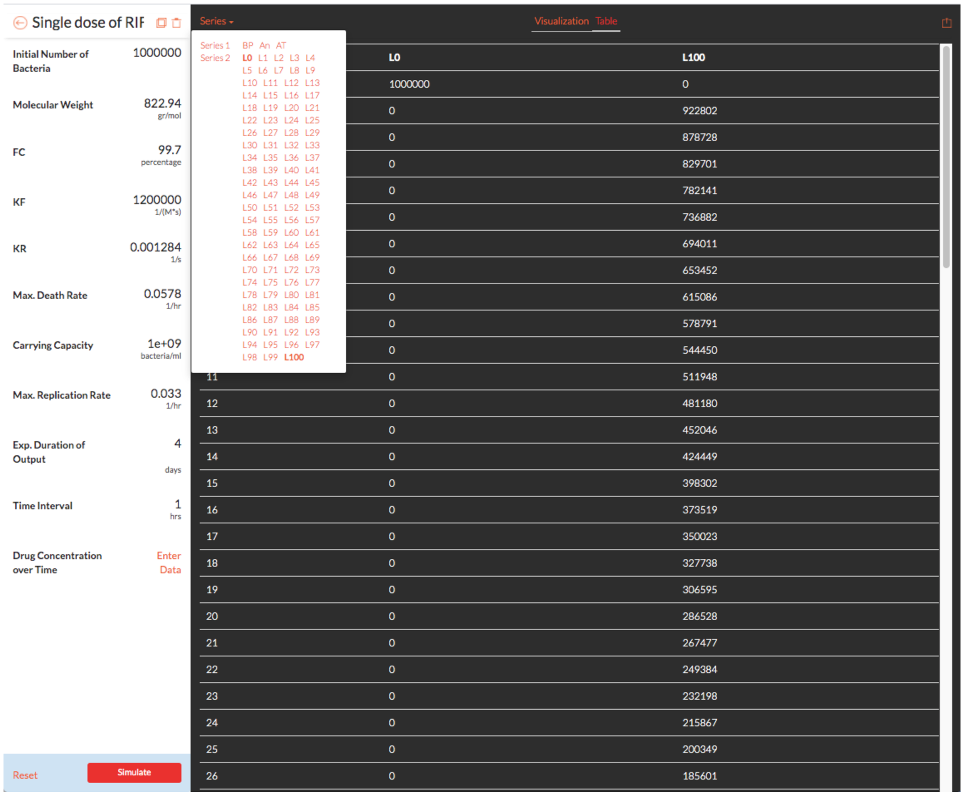


Figure 13: Choose individual data

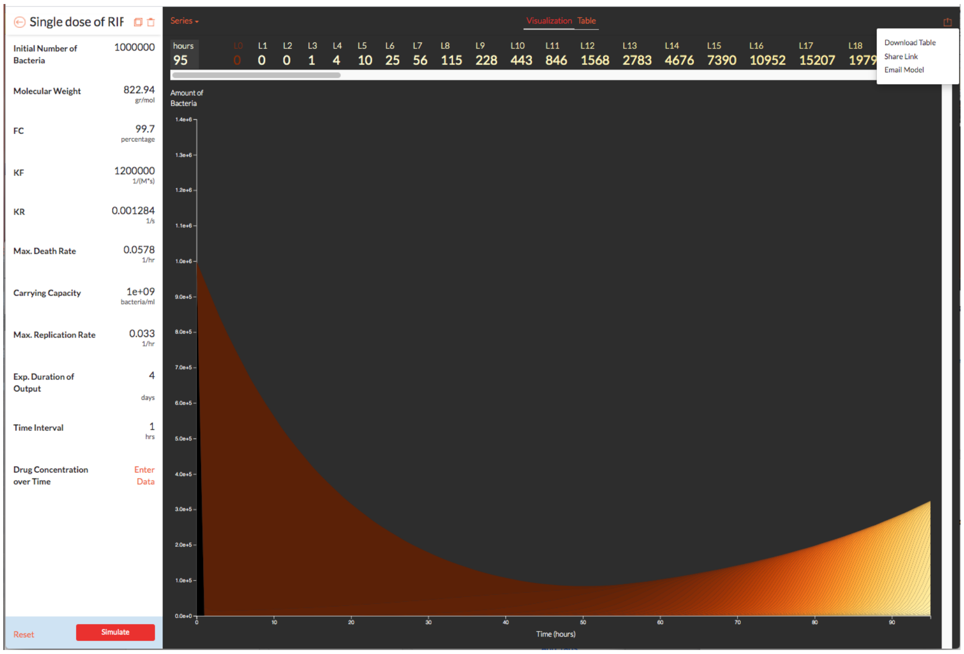


Figure 14: Save and share options

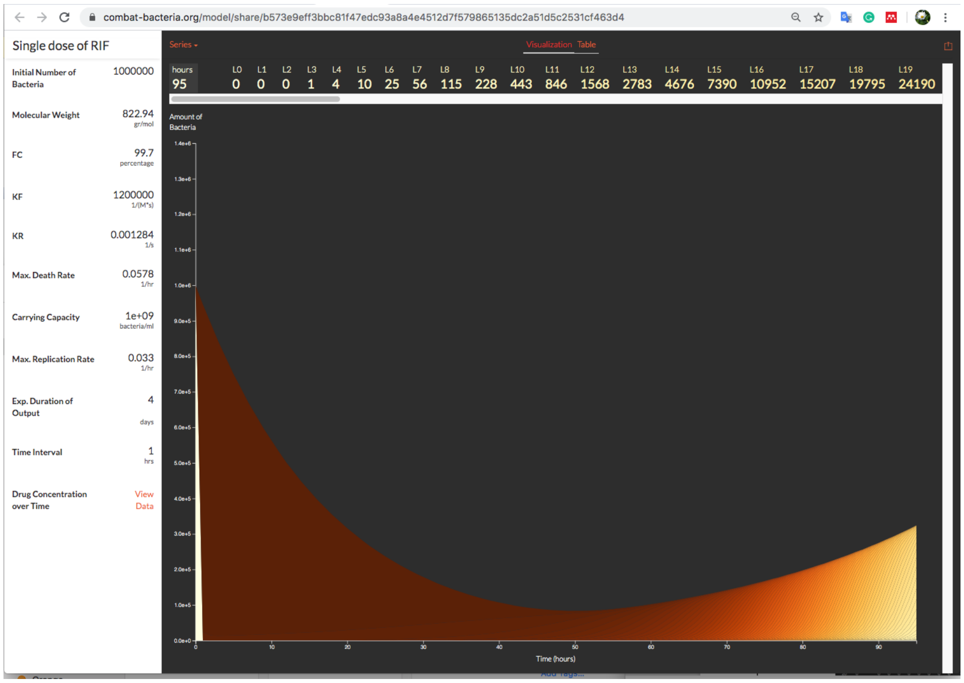


Figure 15: Share models
